# A statistical 3D model of the human cortical vasculature to compute the hemodynamic fingerprint of the BOLD fMRI signal

**DOI:** 10.1101/2020.10.05.326512

**Authors:** Mario Gilberto Báez-Yáñez, Jeroen C.W. Siero, Natalia Petridou

**Author notes:** Correspondence to: Mario G. Báez-Yáñez, PhD Department of Radiology, Imaging Division, University Medical Center Utrecht, Room Q.02.4.307, Heidelberglaan 100, 3584 CX, Utrecht, Netherlands.

## Abstract

BOLD fMRI is a commonly used technique to map brain activity; nevertheless, BOLD fMRI is an indirect measurement of brain function triggered by neurometabolic and neurovascular coupling. Hence, the origin of the BOLD fMRI signal is quite complex, and the signal formation depends, among others, on the geometry of the cortical vasculature and the associated hemodynamic behavior. To characterize and quantify the hemodynamic contributions to the BOLD signal response in humans, it is necessary to adopt a computational model that resembles the human cortical vascular architecture and mimics realistic hemodynamic changes. To this end, we have developed a statistically defined 3D vascular model that resembles the human cortical vasculature. Using this model, we simulated hemodynamic changes triggered by a neuronal activation and local magnetic field disturbances created by the vascular topology and the blood oxygenation changes. The proposed model considers also the biophysical interactions and the intrinsic magnetic properties of the nearby tissue in order to compute a *dynamic* BOLD fMRI signal response. This computational pipeline results in an integrated biophysical model that can provide a better insight on the understanding and quantification of the hemodynamic fingerprint of the BOLD fMRI signal evolution.

## 1. INTRODUCTION

Blood oxygenation level-dependent functional magnetic resonance imaging (BOLD fMRI) is a powerful noninvasive tool which employs susceptibility-sensitive imaging techniques to map in-vivo brain function (Ogawa et al, 1993). Due to the recent advances on hardware at ultra-high magnetic fields (≥ 7T) and the development of new MR pulse sequences, this imaging technique enables the study of brain function at a high level of detail. (Pohmann et al, 2016; Huber et al, 2018; Budde et al, 2014; Shajan et al, 2014; Koopmans et al, 2011; Polimeni et al, 2018; Siero et al, 2013).

However, the BOLD fMRI signal is only an indirect measurement of neuronal function that reflects hemodynamic changes triggered by neurometabolic and neurovascular coupling, and the interaction of the underlying biophysical effects of blood oxygenation and tissue (Buxton et al, 1998; Logothetis et al, 2001; Bandettini et al, 1994).

In order to disentangle the complexity of the BOLD fMRI signal, biophysical and computational modeling of the associated hemodynamic response to a neuronal drive has been a fundamental area of research (Zheng et al, 2002; Zheng et al, 2005; Havlicek et al, 2017; Havlicek et al, 2015; Angleys et al, 2018; Fang et al, 2008; Drysdale et al, 2010).

Several mathematical models exist that characterize the impact of the cerebral blood flow, cerebral blood volume, and oxygen consumption, as they act together in order to sustain the metabolic demand of a neuronal activation or baseline states. For example, the “balloon model” proposed by Buxton et al (1998) and Mandeville et al (1999) or the “vascular anatomical network (VAN) model” of Boas et al (2008).

Nonetheless, it remains unclear how the vascular topology and associated hemodynamics, and their biophysical interaction with nearby tissue, influence the spatial and temporal properties of the BOLD fMRI signal evolution (Petridou et al, 2017; Polimeni et al, 2010; Siero et al, 2011; Barth et al, 2007; Uludag et al, 2018; Buxton, 2012; Markuerkiaga et al, 2016).

Towards understanding the hemodynamic fingerprint of the BOLD fMRI signal formation, computational modeling provides a comprehensive perspective on the relation between the cerebral vasculature and the associated hemodynamic changes (Su et al, 2012; El-Bouri et al, 2015; Payne and El-Bouri, 2018; Linninger et al, 2013). Furthermore, computational approaches offer valuable information to investigate the biophysical interactions between magnetized water molecules that freely diffuse within the tissue and the magnetic susceptibility-induced effects produced by blood oxygenation changes in the vasculature at the mesoscopic level (Kiselev and Posse, 1999). In addition, this modeling approach provides the opportunity to explore and characterize the impact of the MR pulse sequence on the measured BOLD fMRI signal response (Yablonskiy et al, 1994; Uludag et al, 2009).

So far, numerical simulations of the BOLD fMRI signal using non-realistic vascular representations, such as spheres or cylinders, have described *static* signal characteristics. For instance, the influence of the fractional blood volume or the role of the MR pulse sequence dependent on specific parameters of blood and tissue in active and resting states. (Ogawa et al, 1993; Bandettini et al, 1994; Uludag et al, 2009; Boxerman et al, 1995; Weisskoff et al, 1994; Fujita, 2001; Bieri et al, 2007; Scheffler et al, 2019; Markuerkiaga et al, 2016).

Recently, computational simulations using realistic 3D vascular models obtained from the parietal cortex of mice, acquired with two-photon microscopy, have demonstrated that the vascular architecture and the vessel/cortex orientation with respect to the main magnetic field have a considerable influence on the amplitude of the measured BOLD fMRI signal (Gagnon et al, 2015; Báez-Yánez et al, 2017). The latter has also been observed in experimental data (Fracasso et al, 2018; Viessmann et al, 2019).

Rodent models, however, may not be directly applicable to humans due to differences in vascular architecture between species. The geometrical structure of the vascular network between humans and mice is similar but only to a certain degree. For example, the microvasculature (i.e. capillary bed) of humans and mice are correlated through a scaling law (Smith et al, 2019). On the other hand, topological studies of the human cortical vasculature suggest that several human features are not accounted in those realistic 3D mice models (Lorthois et al, 2011). For instance, a significant difference appears on the artery/vein ratio that feed/drain a specific volumetric region that is a ratio of 3:1 in humans as opposed to 1:3 in mice (Uludag et al, 2018; Lorthois et al, 2011; Blinder et al, 2013). These discrepancies might lead to quantitative errors on the computed BOLD fMRI signal and thus to a misinterpretation of data acquired in humans.

At present, a 3D visualization of an ex-vivo human vascular network sample is possible to obtain with two-photon microscopy or x-ray microtomography imaging techniques. However, these technologies are still under development and not trivial to apply, particularly, for sufficiently large tissue samples. Thus, acquisition of large volumes of human cortical vasculature (> 1 mm^3^) is hard to obtain (Cassot et al, 2006; Lauwers et al 2008; Duvernoy et al, 1981).

To circumvent the lack of a realistic 3D representation of the human cortical vasculature, we have developed a computational pipeline that results in a statistical 3D vascular model that approximates the geometrical, topological and rheological characteristics of the human cortical vasculature. In addition, the proposed computational model allows to mimic the temporal evolution of hemodynamic changes in a typical voxel (1 mm^3^) used in high-resolution fMRI, and to simulate intrinsic biophysical properties such as thermal motion of the water molecules (diffusion) and inhomogeneities in the local magnetic field induced by the blood oxygenation variations on the tissue.

The proposed computational pipeline gives an integrated biophysical model that helps to disentangle the underlying anatomical, functional, and biophysical characteristics that constitute the hemodynamic fingerprint of the BOLD fMRI signal, and can give a better understanding of their influence on the spatial and temporal properties of the BOLD fMRI signal evolution.

## 2. MATERIAL AND METHODS

We have developed a computational pipeline that results in an integrated biophysical model to simulate a dynamic BOLD fMRI signal response. The computational pipeline consists of three stages: 1) generation of a statistical 3D vascular model (SVM) of the human cortical vasculature, 2) hemodynamic simulation that occurs in the vascular network given a neuronal activation, and 3) simulation of the biophysical effects induced by hemodynamics and intrinsic magnetic properties of the tissue (diffusion and induced-susceptibility) and the MR pulse sequence.

### 2.1 Generation of a statistical 3D vascular model - SVM

We have developed an algorithm that produces a statistical 3D vascular model of the human cortical vasculature (see Figure 1.f). A representative statistical 3D vascular model is generated in two phases. Each phase approximates a human cortical vessel subpopulation, i.e. a microvascular and a macrovascular network. Finally, these two networks are superimposed to yield a representative human statistical 3D vascular model (SVM). The generation of each vessel subpopulation is described below. Note that in our model, the microvascular and macrovascular compartments are independent structures.

**Figure 1.**
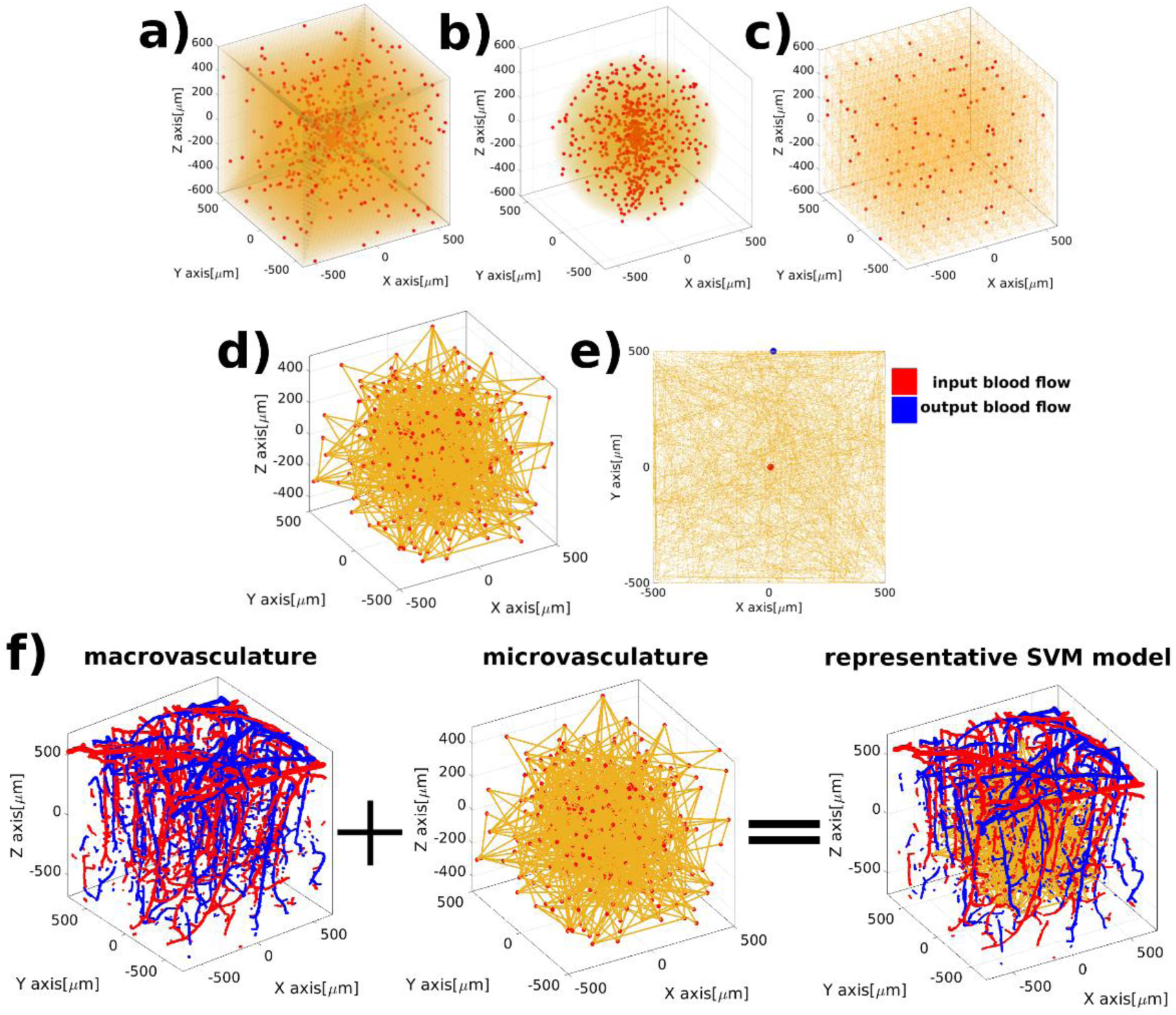
Representative statistical 3D vascular model - SVM. Distribution of random points (vessel joints) in an isotropic space grid composed of cubes (a) spheres (b) and triangles (c). d) pseudorandom connections between vessel joints that approximate statistical features of the microvasculature in humans. e) Representation of the input and output blood source in order to compute the hemodynamics across the network. f) representative 3D SVM. The spatial distribution of the macrovessels (arteries in red and veins in blue) was adapted from the mouse model acquired by Blinder et al (2013). The macrovascular compartment is mathematically modified to reflect the artery/vein ratio obtained from anatomical studies in human cortex (Duvernoy et al 1981).

#### 2.1.1 Generation of the microvascular network

Microvessels are represented as a statistical network that approximates the anatomical properties of the human capillary bed. Properties accounted in the microvascular network generation are the radius of the vessel, the length of the vessel, and the number of connections between vessels. These values are defined according to histological data (Cassot et al, 2006; Lauwers et al 2008; Duvernoy et al, 1981, see Table 1). Moreover, the generation of the microvascular compartment is also limited by a restricted volume fraction. We imposed a ∼3% volume fraction of the total simulation space at basal state.

**Table 1.** Simulation parameters used for a representative statistical vascular model

| architecture parameters |  | hemodynamic parameters |  | biophysical and MR pulse sequence parameters |  |
| --- | --- | --- | --- | --- | --- |
| vessel radius | microvessel<br>Gaussian distr.<br>mean=6.1μm<br>s.d.=1.2μm)<br>macrovessel<br>$\geq 30\mu$m | arterial blood pressure | 75 mmHg | diffusion coefficient tissue | 1μm<sup>2</sup>/ms |
| vessel length | microvessel<br>~10μm-70μm<br>macrovessel<br>100 μm-1000 μm | vein blood pressure | 25 mmHg | static magnetic field strength | 7 tesla |
| number of connections per vessel | ~3-5 connections | partial pressure O<sub>2</sub> in tissue | 15 mmHg | number of spins | 1x10<sup>10</sup> |
| artery/vein ratio | 3:1 | arterial oxygenation level | 90% | echo-time, gradient echo | 27 ms |
| volume fraction microvessel | ~3% | hematocrit | 40% | echo-time, spin echo | 48 ms |
| volume fraction macrovessel | ~2% | arterial dilation | 80%, Gaussian shape, time-to-peak = | time-step | 500 μs |
|  |  |  | 6 s<br>width = 2 s |  |  |
|  |  | Simulation time | 20 seconds | T1 gray matter<br>T2 gray matter<br>at 7 tesla | 1800 ms<br>27 ms |

The microvascular network is generated from the scattering of randomly distributed points inside an isotropic space that mimics the MR imaging voxel (see Figure 1.a, 1.b and 1.c). We constrained the size of the simulation space to 1 mm^3^, which is on the order of a typical voxel size used for high-resolution fMRI. Further, iterative pseudorandom connections between these scatter points are created (see Figure 1.d) until they approximate the statistical density distributions of the human microvasculature according to histological density distributions obtained from experimental data (Smith et al, 2019; Cassot et al, 2006).

To characterize the statistical and geometrical properties of the network, a set of vectors and an adjacency matrix describe the complete structure of the microvascular architecture.

An *m* x 3 vector describes the spatial coordinates of the microvascular vessel joints inside the voxel space (i.e. randomly distributed points described above). The connectivity of the microvascular network is described by an *m* x *m* microvessel connectivity matrix M, i.e. an adjacency matrix, where each element M*_ij_* is equal to one if the node *i* is connected to the node *j*, and zero otherwise.

Moreover, each microvessel (physical connection between vessel joints) is simulated as a finite cylinder. Spatial information of each microvessel (cylinder) is stored in an *n* x 6 information vector (spatial position and direction of the cylinder).

#### 2.1.2 Superposition of the macrovascular architecture

Pial and penetrating arteries and pial and ascending veins comprise in our model the macrovascular compartment (see Figure 1.f). Arteries and veins were defined based on the 3D vascular architecture obtained from the parietal cortex of mice acquired by Blinder et al (2013) with two-photon microscopy. The mouse macrovascular compartment was segmented based on a vessel radius threshold. Vessels larger than 30 µm were selected and defined as the macrovascular compartment.

Further, the mouse macrovascular compartment was mathematically modified to approach human brain characteristics, according to the artery/vein ratio in humans of 3:1 (Duvernoy et al, 1981) as opposed to 1:3 in mice (Uludag et al, 2018; Blinder et al, 2013). The modified macrovascular compartment consists of ∼2% of the total volume fraction in the simulation space.

Finally, the modified macrovasculature is superimposed on the microvascular compartment as depicted in Figure 1.f to yield a representative statistical 3D vascular model.

To characterize the topology of the macrovascular network, similarly as with the microvasculature, the spatial position corresponding to each artery or vein segment, simulated as a finite cylinder, is stored in a *p* x 6 information vector (spatial position and direction of the cylinder).

### 2.2 Hemodynamic simulation using the SVM

Hemodynamics are simulated based on the principle of conservation of mass and energy. We compute the blood flow, blood pressure and blood oxygenation changes considering the rheological properties of blood and anatomical features of the vessels.

It should be mentioned that the computation of the hemodynamic changes in the microvascular and macrovascular compartments in our model are independent. The microvascular compartment presents a dynamic blood evolution, while the macrovascular compartment presents a pseudo-dynamic blood behavior.

#### 2.2.1 Simulation of microvascular hemodynamics

Due to the mathematical similarities between an electric circuit and fluid dynamics, the statistical microvascular network is translated into an electric circuit representation to simulate the evolution of hemodynamics in the network (Boas et al, 2008). The hemodynamic simulation gives as a result the evolution of blood flow, blood pressure and blood oxygenation behavior across the microvascular network.

We considered that the microvessel resistance (R) is described by the Poiseuille’s law

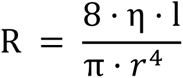

that depends on the radius of the microvessel (r), the length of the microvessel (l) and the viscosity of blood (η). Further, the hemodynamic simulation takes into account the rheological properties of blood flow, specifically the Fahraeus and the Fahraeus-Lindquist effects that describes how the viscosity of a fluid, in this case blood, depends on the radius of the microvessel and the hematocrit level (Su et al, 2012; Pries et al, 1990).

We imposed a constant level of hematocrit (Htc) across the whole microvascular network (Htc = 40%). At present, we assumed that all microvessels present the same structural elements, that is, we do not take into account the influence that vascular muscle cell type or pericytes will have on the resistance behavior of each particular microvessel or their impact on the blood flow control.

The microvasculature is fed by only one artery (the input blood flow source) and is drained by one vein (the output blood flow). The spatial position of the input/output blood sources can be chosen arbitrarily across the network. Here, we established that the artery is positioned in the center of the microvascular network and the vein is randomly positioned in an outer vessel joint of the network (see Figure 1.e). The artery was set to present a blood pressure of 75 mmHg, and the vein that drains the blood out of the network was set to a blood pressure of 25 mmHg at basal state (Vovenko, 1999).

The computation of a basal blood flow and blood pressure state in each microvessel across the network is straightforward to compute based on the conservation of mass

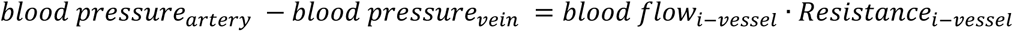

which describe the relation between the blood pressure in the inlet/outlet source point and the microvascular resistance properties as described above (Boas et al, 2008; see Table 1).

#### 2.2.2 Macrovascular compartment: pseudo-dynamic blood behavior

The macrovascular compartment is assumed to present pseudo-dynamic blood properties. The properties simulated on the input (one artery) and output (one vein) sources of the microvascular network are used to drive the blood oxygenation level of the entire arterial and venous macrovascular compartment for each time-step.

For the arterial compartment, we assumed that the blood oxygenation level is constant over time for all arteries (95% oxygenation level) given the fact that arteries do not present a large change in blood oxygenation upon neuronal activation. The impact of volume changes on the computed signal is discussed below.

For the venous compartment, the blood oxygenation level is determined by the differential blood oxygenation calculated at the output source (i.e. the draining vein) for each time-step from the hemodynamic simulation of the microvascular compartment, thus the pseudo-dynamic behavior for the macrovascular compartment.

#### 2.2.3 Hemodynamic simulation as a result of a global neuronal activation

Hemodynamic changes resulting from neuronal activation are computed as the differential change of the blood flow, pressure, and oxygenation level in each microvascular segment.

In order to calculate the oxygen saturation fraction of each microvessel, the oxygen content in a vascular segment is a balance between oxygen flow into the i-th microvessel via blood flow and out of the i-th microvessel via blood flow and oxygen diffusion into the tissue, as scheme in the next relation,

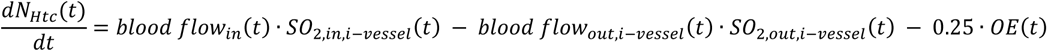

where *SO_2,in,i−vessel_* and *SO_2,out,i−vessel_* is the oxygen saturation flowing in and out of the i-th vessel respectively. OE is the oxygen extraction fraction from the i-th vascular segment into the tissue. For a detailed description of the oxygen dynamics, see (Boas et al, 2008).

It is considered that the tissue maintains a cerebral metabolic rate of oxygen (CMRO_2_) inside the simulation space that depends on its partial pressure of oxygen. We assumed a partial pressure of oxygen in tissue of 15 mmHg (Vovenko, 1999).

For simplicity, we assumed that the tissue consumes exactly the same amount of oxygen independently of the spatial location of each microvessel segment, i.e. we do not simulate a spatially dependent neuronal activation, but a global neuronal activation (Secomb et al, 2004).

Thus assuming a global neural activation, an arterial dilation is simulated as a blood flow increase at the input point to satisfy the demand of the neurovascular coupling. This results in a differential hemodynamic transition state for each vascular segment across the microvascular network (Huppert et al, 2007).

The arterial dilation, which effectively changes arterial resistance, is modeled as a Gaussian function with properties of time to peak at 6 seconds and a width of 2 seconds. For illustration purposes we simulated an 80% arterial dilation as compared to basal state to emphasize the hemodynamic changes predicted from the model (see Figure 3 and 4). A hemodynamic simulation with physiologically meaningful arterial dilation parameters (10%) can be found in the Supplementary Data.

**Figure 2.**
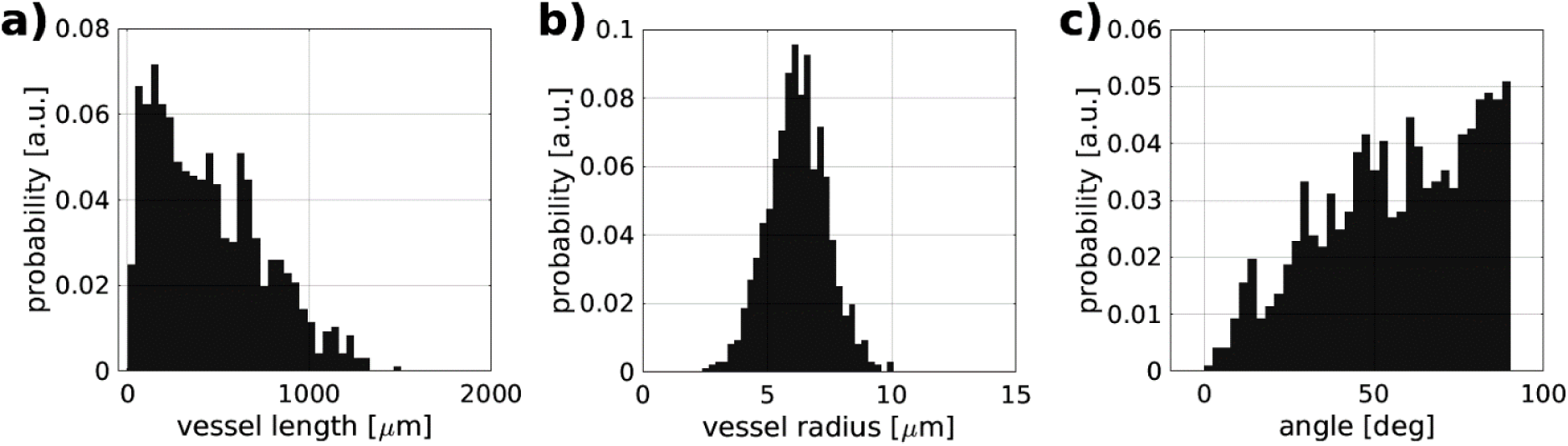
Statistical description of a representative microvascular compartment: density distribution of the vessel’s length (a), of the vessel’s radius (b), and of the vessel’s orientation with respect to the cortical surface (c).

**Figure 3.**
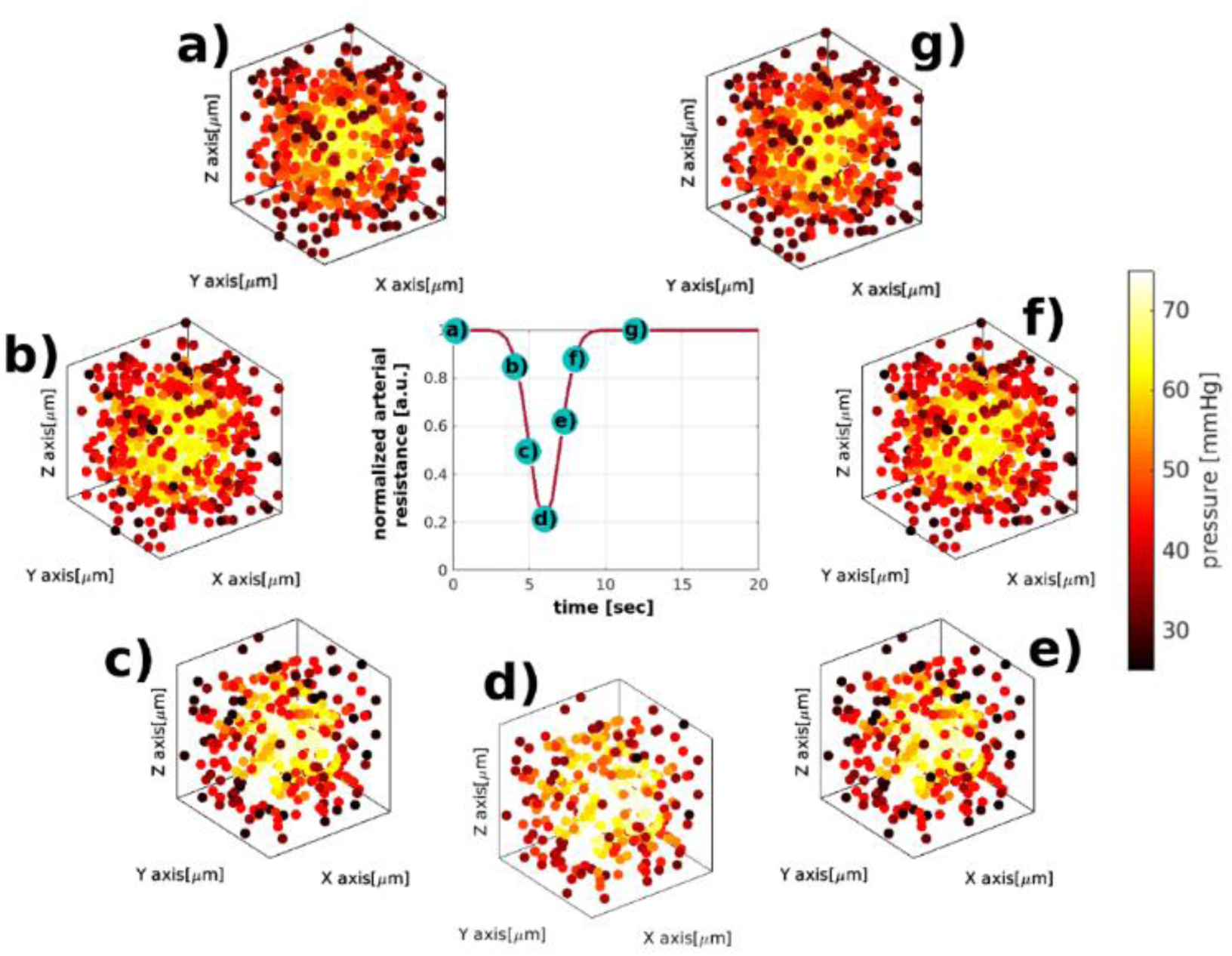
A representative evolution of hemodynamics on the microvasculature for an arterial dilation of 80%. For illustration purposes, only the vessel joints are displayed. One can observe an increase of the blood pressure across the network in a radial-like shape as an effect of the arterial dilation and conservation of mass across the network.

**Figure 4.**
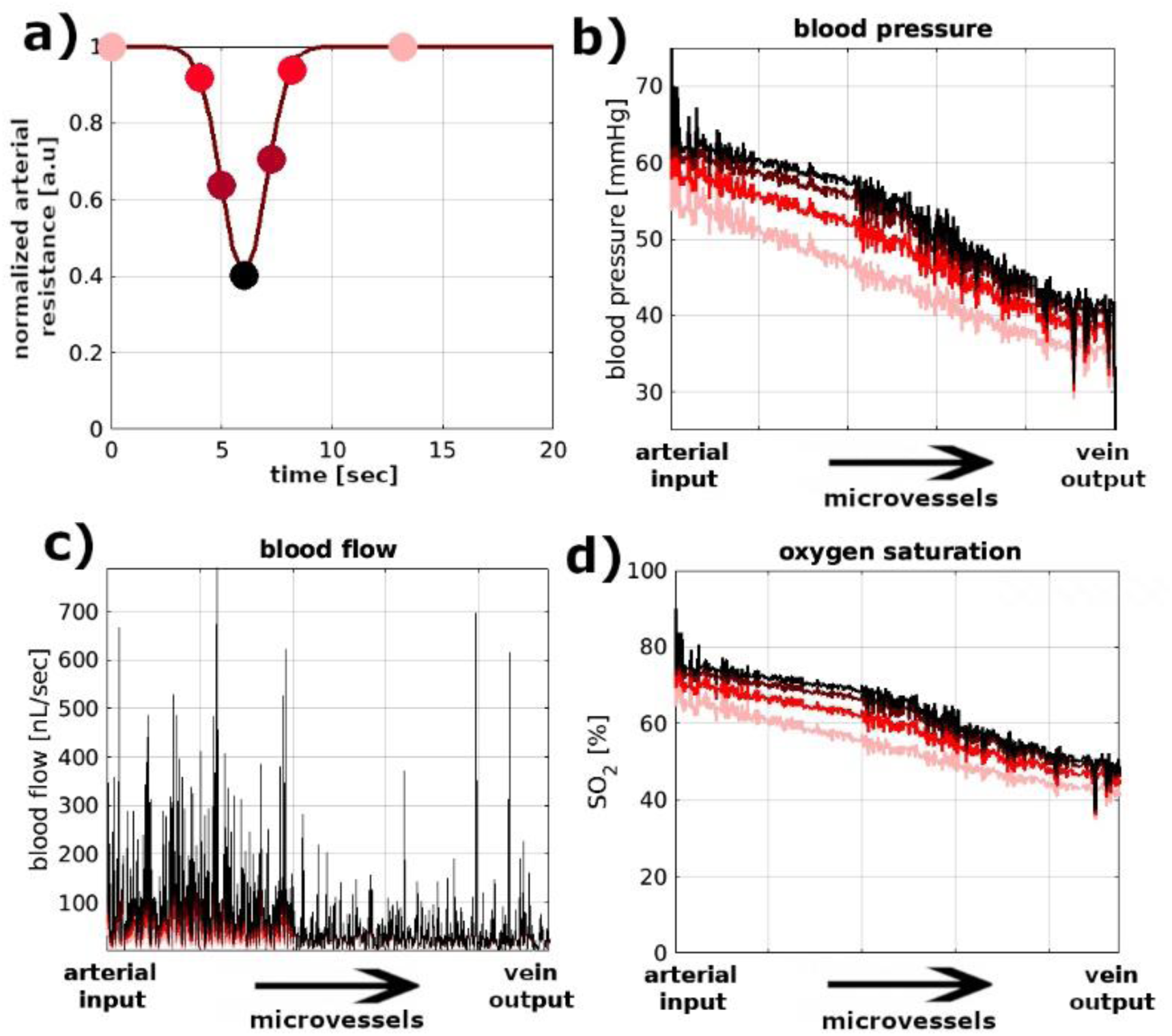
Hemodynamic response of the microvascular network to a differential change in the arterial resistance (a). Four curves are plotted for different time points of the arterial dilation indicated by the colored circles in panel (a): an increase in blood pressure (b), blood flow (c), and blood oxygenation level (d). The black arrow denotes a linear representation of the 3D microvascular network between the artery (input blood source) and vein (output blood source).

### 2.3 Simulation of the biophysical effects induced by hemodynamics and intrinsic magnetic properties of the tissue (diffusion and induced-susceptibility) and the MR pulse sequence

Given each differential hemodynamic stage over time (time-step of 0.5 ms), it is possible to compute a differential local magnetic field for each time-step. For the calculations, we assumed the dipolar magnetic response of a finite cylinder, i.e. vessel strand, with neglected boundary conditions on the extremities for both the microvascular and macrovascular compartment (Báez-Yánez et al, 2017).

The spatial information vectors of each microvessel and macrovessel segment are used to compute a local magnetic field disturbance in the simulation space for each time-step using the differential blood oxygenation level of each vessel obtained from the hemodynamic simulation (section 2.2.3).

It is important to mention that even though the macrovascular compartment presents a pseudo-dynamic blood behavior, the geometry and the differential blood oxygenation level of macrovessels contribute to the local magnetic field inhomogeneity at each time-step, and thus then to the computed BOLD signal response. With this approach, we simulate similar conditions as compared with other models (Uludag et al, 2009; Scheffler et al, 2019; Markuerkiaga et al, 2016), but we go further by adding the dynamics of the microvascular compartment.

In addition, an ensemble of magnetized water molecules, or spins, depicting thermal motion is simulated through a Monte Carlo approach. The ensemble of molecules senses the extravascular local magnetic field disturbances with a diffusion coefficient of 1 µm^2^/ms. We considered that the exchange of water molecules between vascular compartments is prohibited, i.e. we assume an impermeable vascular network.

In order to contain all spins inside the simulation space, voxel boundary conditions are established as an infinite space, i.e. if a spin moves out of the voxel, it is then mirrored on the opposite side of the simulation space preserving the magnetization history.

A dynamic BOLD fMRI signal can be computed at any magnetic field strength considering the proper relaxation times of surrounding tissue and blood. We used the non-linear relation of the longitudinal and transversal relaxation times with respect to the magnetic field strength given by Khajehim et al (2017) for the gray matter at 7T (see T1 and T2 values in Table 1).

The computational pipeline allows positioning of the main magnetic field in any spatial direction. We define the angular orientation (φ = 0 degrees) of the SVM to be parallel between the main magnetic field (z-axis) and the normal vector that rises from the cortical surface. The cortical surface does not present any curvature along the simulation space, i.e. pial arteries and veins are orthogonal to the main magnetic field (Fracasso et al, 2018; Viessmann et al, 2019).

In addition, to demonstrate the impact of the angular orientation of the human vasculature with respect to the main magnetic field on the amplitude of the BOLD signal, we computed the local magnetic field variations of two exemplary time-points, i.e. a representative active and resting state. The inhomogeneous magnetic field calculation assumes in the active state an oxygen saturation in the arteries of 98%, microvessels of 85% and veins of 75%; and in the resting state the vasculature present an oxygenation saturation of 95% - 75% - 68% in the vascular compartments, respectively. The MR signal evolution for both states is calculated as the Fourier transform of the computed local frequency components in each state (see Figure 5, Ziener et al, 2007). This kind of calculation assumes a gradient-echo pulse sequence and no diffusion effects, i.e. static diffusion regime.

**Figure 5.**
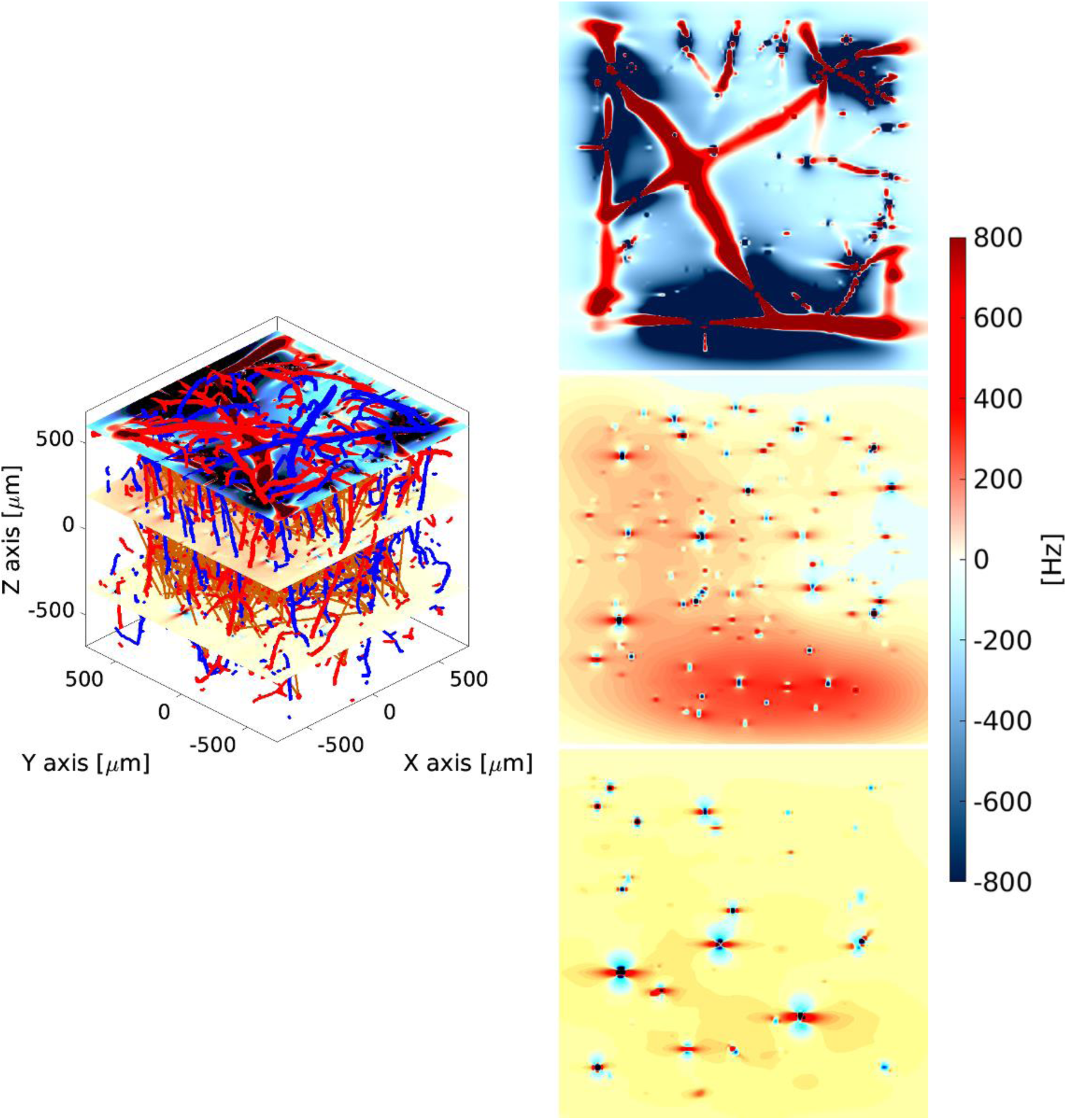
Inhomogeneous magnetic field produced by a representative SVM for an exemplary hemodynamic time-point. Magnetic field maps are shown for three different locations inside the vascular model: top, middle and bottom planes. The magnetic field maps represent parallel planes with respect to the cortical surface (xy-plane). The main magnetic field lies on the z-axis, i.e. orthogonal to the cortical surface.

To emphasize the effect of a particular vascular compartment, we compute the local magnetic field variation for: I + II) pial arteries and pial veins plus penetrating arteries and diving veins (complete macrovasculature), II) only penetrating arteries and diving veins, III) capillary bed (microvasculature), I + II + III) complete SVM model, and finally, II + III) penetrating arteries and diving veins plus microvasculature (see Figure 7).

**Figure 6.**
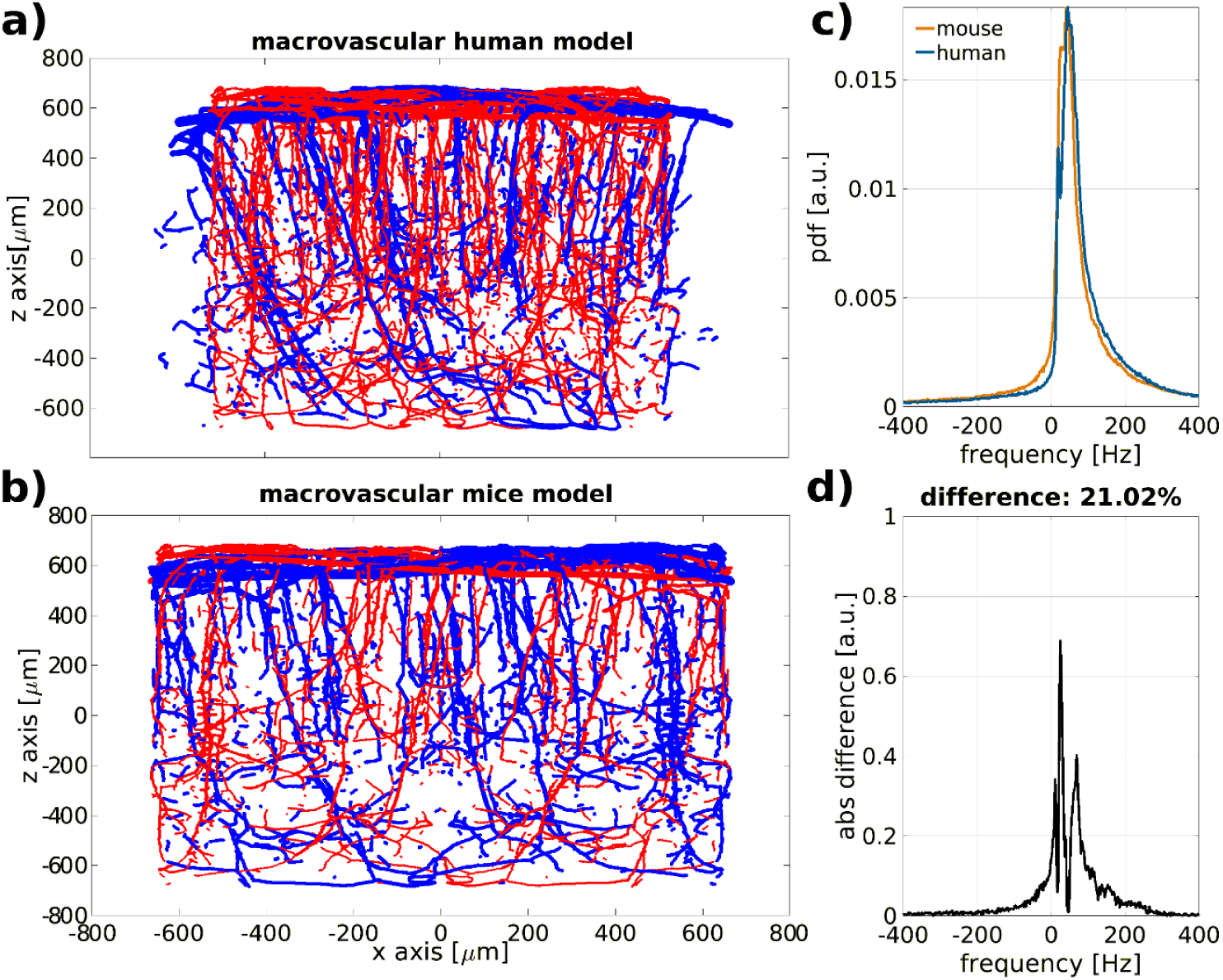
Representative macrovascular architecture between human (simulated data, panel (a)) and mouse (experimental data, panel (b)) and frequency line-shape comparison (panels c and d). The arterial compartment is shown in red and the venous compartment in blue, c) frequency line-shape computed at 7T for each model and d) frequency line-shape difference between human and mouse macrovascular compartment. There is a ∼21% difference in the inhomogeneous magnetic field between the human and mouse model for the same blood properties. (arteries = 95% oxygenation level, veins = 65% oxygenation level, Htc = 40%).

**Figure 7.**
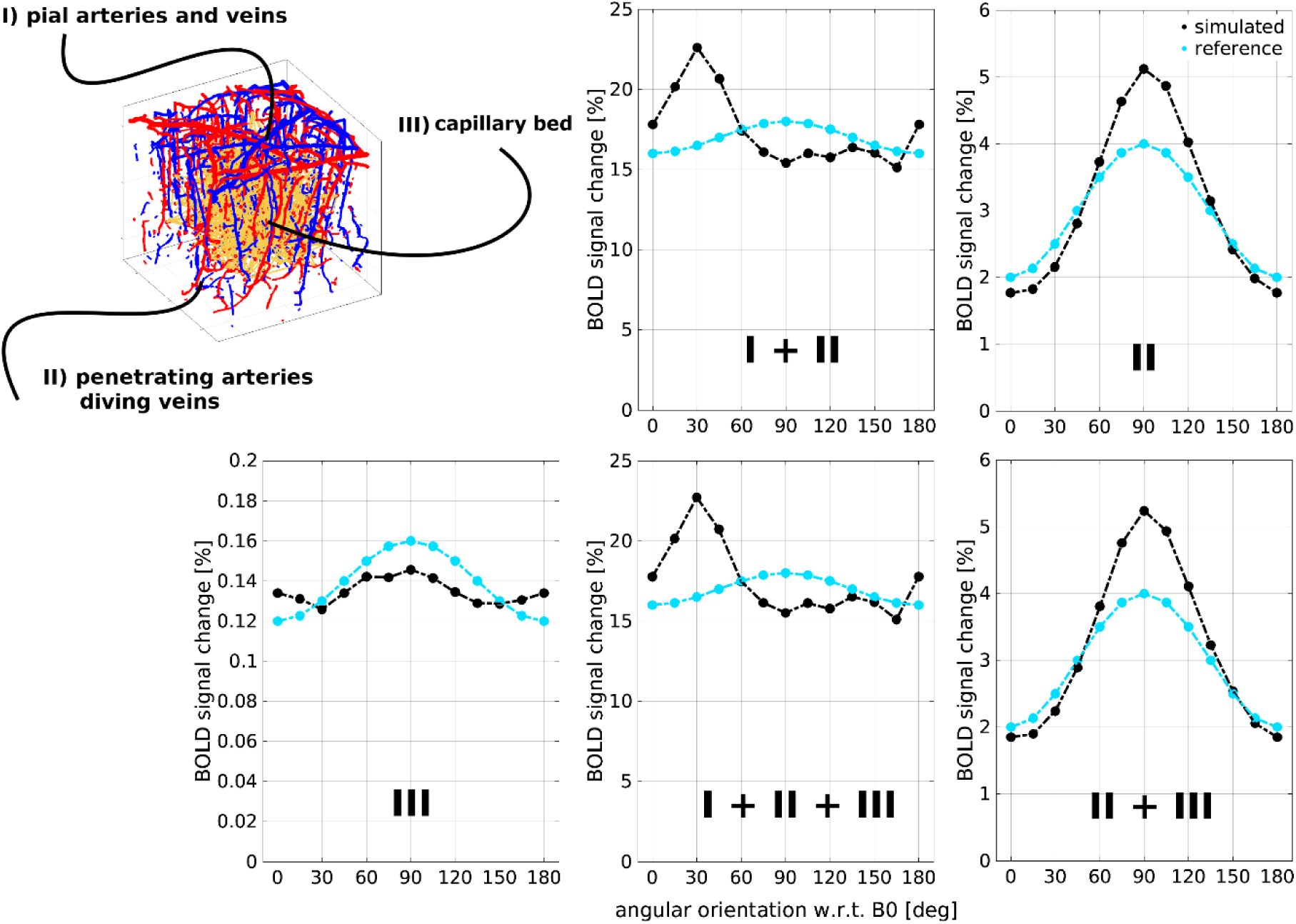
Impact of the angular orientation of the simulated human vasculature on the BOLD signal amplitude. We compute the local magnetic field variation for I + II) pial arteries and pial veins plus penetrating arteries and diving veins (i.e. complete macrovasculature), II) only penetrating arteries and diving veins, III) microvasculature, I + II + III) complete SVM model, and II + III) penetrating arteries and diving veins plus microvasculature. The x-axis of all plots represents the angular orientation (φ) between the main magnetic field and the normal vector that rises from the cortical surface. Note that the scale of the BOLD signal change (y-axis) is notably different for the different vascular combinations. The light-blue line represents a scaled reference to facilitate the visualization of the cos^2^(φ) obtained in the experimental data (Fracasso et al, 2017; Viesmann et al, 2019).

Finally, the BOLD fMRI signal is computed by the interaction of the moving spins within each differential local inhomogeneous magnetic field (Boxerman et al, 1995). The dephasing of all spins is integrated over the echo-time (TE) corresponding to a particular differential hemodynamic stage.

Simulations shown here were run for a bulk of 1×10^10^ spins for gradient-echo (TE = 27 ms) and spin-echo (TE = 48 ms) at 7T with a time resolution of 500 µs for a total response time of 20 seconds with an angular orientation of the normal vector of the SVM parallel to main magnetic field (φ = 0 degrees). The bulk of spins depict motion in a spatial continuum space. The simulation pipeline is implemented in Matlab/Julia and was computed on a CPU cluster server with 40 cores. Matlab codes to simulate the SVM model in this study are available upon request. All simulation parameters used to compute the representative dynamic BOLD fMRI signal are summarized on Table 1.

## 3. RESULTS

### 3.1 Generation of a statistical 3D vascular model

From Figure 1.a to Figure 1.c, we display three possible configurations that can be used to create the microvascular compartment. The main difference between these configurations is the way of scattering the random points (vessel joints) contingent on convergence into a microvascular representation that statistically satisfies the properties of the human microvasculature. It is clearly visible that all three configurations show different spatial arrangements. This property allows to simulate different microvascular geometries within a voxel.

Figure 1.d illustrates the pseudorandom connections between the distributed points that constitute the statistical description of the microvascular network. Using vector analysis and graph theory (section 2.1.1), it is straightforward to describe the properties of the microvascular compartment as shown in the density distributions in Figure 2.

The iterative pseudorandom generation of connections assures convergence to a statistical description of the microvascular network that approximates properties reported in humans (Cassot et al, 2006; Lorthois et al, 2011). Then, the mathematically modified macrovascular compartment is superimposed on the microvascular network (Figure 1.f) resulting in a representative statistical 3D vascular model - SVM.

The representative SVM depicted on Figure 1.f consists of 713 microvessels, 319 closed loops, a ∼5% volume fraction occupied by all vessels and an artery/vein ratio of 3:1.

As observed in Figure 2.b, we imposed on the vessel radius for the microvascular compartment a Gaussian distribution with a mean radius of 6.1 µm and a standard deviation of 1.3 µm. The most occurring vessel length is ∼50–60 µm. The vessel size of the macrovascular compartment was defined by the experimental data obtained by Blinder et al, (2013) and consist of vessels with radius sizes within the range of 30 to 70 µm.

### 3.2 Hemodynamic simulation using the SVM

Figure 3 shows the hemodynamic response of the microvascular network given by an arterial dilation, i.e. differential change in the input resistance, located in the center of the simulated voxel (see Figure1.e). Dependent on the arterial dilation state, the blood pressure increases in a radial-like pattern away from the arterial source.

The 3D vascular network is transformed into a linear representation where it is possible to analyze the variation of blood dynamics between the arterial source (input blood flow) and the draining vein (output blood flow).

The hemodynamic changes (blood pressure, blood flow and blood oxygenation level) on the microvascular compartment that result from the arterial dilation are shown in Figure 4. One can observe a sigmoidal-like shape that reflects the decrease of blood pressure and blood flow with respect to distance from the arterial source to the venous output.

The blood oxygenation level per vessel segment is dependent on the blood flow, and given the assumed homogeneous oxygen consumption inside the simulated voxel, it follows the same sigmoidal like-pattern.

### 3.3 Simulation of the biophysical effects induced by hemodynamics and intrinsic magnetic properties of the tissue (diffusion and induced-susceptibility) and the MR pulse sequence

In Figure 5, a representative SVM model is schemed with three magnetic field maps that represent the local inhomogeneous magnetic field produced at the top, middle and bottom planes for an exemplary time-point of the simulated hemodynamic response.

The small circles and dipolar shapes in the middle and the bottom planes represent the contribution of the penetrating arteries and diving veins, respectively. The top plane highlights the magnetic field distortion induced by the macrovascular compartment that lies near the cortical surface. The main distortion of the magnetic field at the top plane comes from the veins and their relative low blood oxygenation level. Microvascular structures are not visually discernable in the maps but their properties contribute to the distortion of the local magnetic field.

The main parameters that alter the differential local magnetic field are the blood oxygenation level of each vascular segment, the vessel radius and its angular orientation with respect to the main magnetic field.

In order to demonstrate the effect of the difference of the artery/vein ratio on the inhomogeneous magnetic field produced by the macrovascular architecture, in Figure 6 we display the macrovascular magnetic field distortion produced by an artery/vein ratio for humans (3:1) and mice (1:3). The computation assumes a static blood condition with 95% oxygenation level for the arterial compartment and a 65% oxygenation level in the veins. We computed a ∼21% difference on the characteristic Larmor frequencies between the simulated inhomogeneous magnetic fields.

To exhibit the impact of the angular orientation of the human vasculature with respect to the main magnetic field on the amplitude of the BOLD signal, we computed the local magnetic field variations of two exemplary time-points, i.e. a representative active and resting state. In Figure 7, we show the angular orientation-dependence of a representative SVM at different orientations with respect to the main magnetic field and a vector normal to the cortical surface (from 0 degrees to 180 degrees at intervals of 15 degrees). The MR signal evolution for both states is calculated as the Fourier transform of the computed local frequency components in each state (see Figure 5, Ziener et al, 2007). To emphasize the effect of a particular vascular compartment, we compute the local magnetic field variation for: I + II) pial arteries and pial veins plus penetrating arteries and diving veins (complete macrovasculature), II) only penetrating arteries and diving veins, III) capillary bed (microvasculature), I + II + III) complete SVM model, and finally, II + III) penetrating arteries and diving veins plus microvasculature.

We can observe that the reported cos^2^(φ) behavior by Gagnon et al (2015), Báez-Yánez et al (2017) and Fracasso et al (2017) is preserved on the simulations where the pial arteries and veins are not accounted in the calculations. On the other hand, the angular behavior when the pial arteries and veins contribute to the local frequency variations does not follow the experimental cos^2^(φ) response. Moreover, the microvasculature presents a negligible angular dependence.

It is also worth noting what the effect of a particular vascular compartment is on the amplitude of the BOLD signal. For instance, the computed %BOLD signal change when the pial vessels are included in the simulation, produce a signal modulating effect that is approximately four times larger compared to when only intracortical vessels contribute to the BOLD signal formation. Microvessels, when isolated, show a relatively small BOLD signal change as compared to the signal change induced by the macrovessels.

Finally, in Figure 8, we show the BOLD signal evolution computed with the Monte Carlo simulation for a gradient-echo and spin-echo pulse sequence. This result is obtained from the complete statistical 3D vascular model, i.e. considering the hemodynamic simulation for the microvascular compartment and the pseudo-dynamic blood behavior for the macrovascular compartment (see section 2.3).

**Figure 8.**
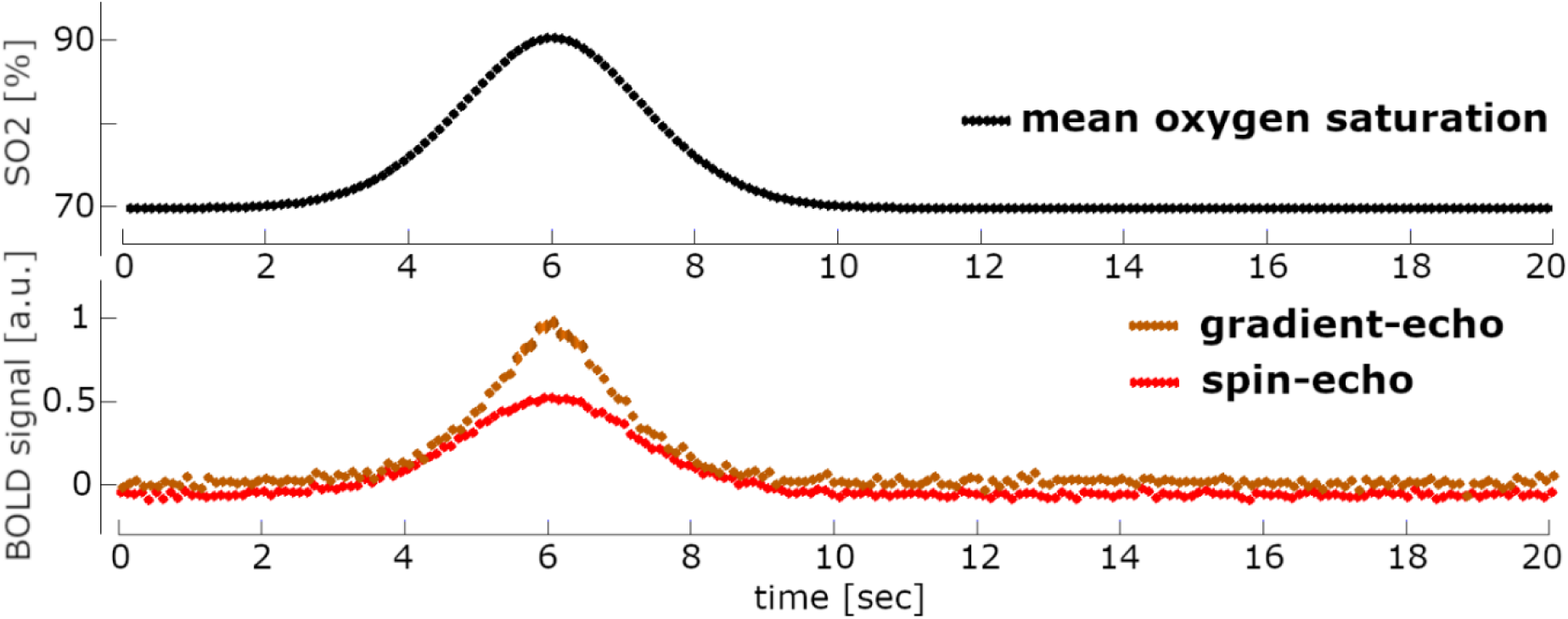
Dynamic BOLD fMRI signal response obtained for an arterial dilation (mean oxygenation level) for a gradient echo (TE = 27ms) and a spin-echo (TE = 45ms) pulse sequence at 7T.

We observe that, as expected, a gradient-echo response is different from a spin-echo response. The main difference between MR pulse sequences lies on the amplitude of the BOLD fMRI signal - proportional to two-fold larger for a gradient-echo acquisition-given by the large contribution of the macrovascular compartment for the gradient-echo sequence.

Given the fact that our model assumes only a linear behavior of the vessel (i.e. vessels mimicked only as a resistance component, see Section 2.2.1), the expected hemodynamic response function (HRF) characteristics such as the onset and the undershoot do not shown any modification in the simulated BOLD signal responses.

## 4. DISCUSSION

To circumvent the current unavailability of a realistic 3D human vascular model, we have developed a novel statistical 3D vascular model that closely approximates the geometrical, topological and rheological characteristics of the human cortical vasculature. The proposed computational model allows to generate different geometrical distributions of vessel subpopulations yielding, for instance, simulations of different microvascular architecture characteristics, such as density and laminar distributions, as may be observed in the human brain. The proposed computational pipeline enables simulation of the temporal evolution of hemodynamic changes in a 3D fashion, and computation of the biophysical effects induced by specific blood oxygenation changes, i.e. the dynamic BOLD fMRI signal response.

### 4.1 On the proposed vascular model, the SVM

We have developed a novel statistical 3D vascular model that closely resembles the geometrical, topological and rheological characteristics of the human cortical vasculature.

The presented vascular model shows advantages compared to other computational approaches. For example, the use of a statistical 3D model gives valuable insights about the direct effect of the vascular morphology and topology on the MR signal formation, in comparison to Markuerkiaga et al (2016) which simulations are computed in a 2D model. Further, the use of a statistical vascular model diminishes the use of non-realistic models which significant assumptions are circumvented, such as the uniform angular distribution of vessels with respect to the main magnetic field and/or the composition of the vasculature defined by a particular vessel radius size (Scheffler et al, 2019; Uludag et al, 2009).

A key advantage of our approach is that it mitigates the need for an experimental 3D reconstruction of the human microvasculature, which is currently difficult to obtain as the microvasculature is statistically defined. Moreover, the ability to generate different geometrical distributions of microvessels enables simulation of several cortical regions with different microvascular topologies as observed in the human brain (Cassot et al, 2006; Lauwers et al 2008; Duvernoy et al, 1981). In addition, the morphology and topology of the macrovascular compartment provide insights on the impact of the vascular architecture on the evolution of the BOLD fMRI signal.

It is important to note that there are few research groups working on the improvement of techniques that can be used to visualize human vascular networks. Most of the human cortical data is acquired with two-photon microscopy imaging or scanning electron microscopy of vascular corrosion casts (Lorthois et al, 2011; Cassot et al, 2006; Duvernoy et al, 1981). Nevertheless, the acquisition of large volumes of such data (∼2 to 4 mm^3^) continues to be an expensive task, time-consuming and it can only be tackled with significant expertise on the field (Blinder et al, 2013; Weber et al, 2008; Gould et al, 2017) limiting the field of view to a few micrometers (∼400 µm^3^). Besides the mice models used by Gagnon et al (2015) and Báez-Yánez et al (2017), to the best of our knowledge, there is no human cortical vasculature reconstruction in a three-dimensional form from a large volume sample (>1mm^3^) at present.

Uludag et al (2018) and Schmid et al (2019) made an extensive analysis across different cortical vascular architectures from different species. An important difference between species lies on the specific volumetric value of blood supplied by the arteries or drained by the veins. They conclude that the macrovascular compartment between mice and humans differs only by the arterial/venous ratio (1:3 in mice versus 3:1 in humans). Yet, macrovascular morphology and topology seems to be conserved across species. This analysis supports the complement of our 3D vascular model with the experimental macrovascular data of Blinder et al (2013) but mathematically modified to fulfill the measured artery/vein human ratio. However, the cortical thickness for humans is larger than that of mice. Nevertheless, we are interested here on the vasculature influence on the MR signal as a whole and not, particularly, in the laminar contribution. Thus, the cortical depth will not modify the computed signal while the arterial/vein ratio keeps constant across a particular simulation space. For investigations of the laminar vascular contributions to the BOLD signal, the cortical depth, the artery/vein ratio and fractional volume can be modified in our model to match different laminar distributions.

Histological studies on ex-vivo samples performed by Lauwers et al (2008) determined that the orientation of the microvascular compartment seems to be not purely random. Nevertheless, most of the non-realistic vascular models assume that the microvasculature is randomly oriented with respect to the cortical surface (Markuerkiaga et al, 2016; Kiselev and Posse, 1999; Uludag et al, 2009; Boxerman et al, 1995). In our approach, the microvascular compartment is generated based on the iterative convergence of the spatial distribution and number of vessel connections to the distributions obtained from histological data, where the angular orientation of vessels is not a defined parameter. This approach did not yield a uniform distribution of the angular orientation while avoiding a considerable violation on the imposed statistical properties (length, number of connections) for any of the three configurations. We might conclude that it is topologically improbable to generate a randomly oriented microvascular compartment that keeps the other statistical features intact and within the experimentally obtained values for the human cortical microvasculature.

Furthermore, Su et al (2012) proposed a microvascular network that serves as a model to calculate blood flow patterns. In our proposed model, we complement the analysis of the statistical 3D vascular model including several other parameters. Specifically, the density distribution of the length of the microvessel and their angular dependence with respect to the cortical surface. These parameters are important to account for an accurate simulation of the hemodynamics (direct impact on the Poiseuille’s law) and the computation of the local inhomogeneous magnetic fields (vessel orientation with respect to the main magnetic field), respectively.

Hence, the proposed vascular model resembles, at least statistically, the geometry and topology of the human cortical microvasculature in a three-dimensional manner where hemodynamics can be simulated. Towards a more accurate characterization of any region of the human brain with our model, a better quantitative and qualitative statistical description of the architecture of the cerebral vasculature is required. These properties are important to compute an adequate disturbance on the magnetic field maps and a reliable evolution of the BOLD fMRI signal.

### 4.2 On the hemodynamics

The proposed computational pipeline allows to simulate hemodynamic changes in a 3D vascular network.

Several models have been proposed to simulate the hemodynamic behavior of cerebral blood flow and blood volume changes in a 2D vascular network and/or with assumptions on the behavior of a particular vascular compartments (Buxton et al, 1998; Havlicek et al, 2015; Mandeville et al, 1999; Boas et al, 2008). The present vascular model accounts for all three compartments –arteries, capillaries and veins-assuming their morphology and spatial distribution described by statistical density distributions. Thus, it offers the possibility to study the hemodynamic behavior in a 3D network and the influence of the spatial contribution of the vasculature to the BOLD fMRI signal behavior. Hemodynamic simulations on the microvascular network are shown for one artery (input blood source) and one vein (output blood source). However, the number of sources (inputs/outputs) can be extended to analyze the evolution of hemodynamics for a different number of arteries or veins.

It is well known that boundary constrains have a strong influence on the computation of the blood flow distribution over the vascular network (Lorthois et al, 2011). For instance, we assumed that the simulation voxel is completely isolated from blood contributions from neighboring voxels. This assumption might influence the results. However, given the fact that the hemodynamic point spread function is limited to several hundreds of micrometers (Vazquez et al, 2014), it can be expected that our model gives a truthful hemodynamic approximation when the input sources lie in the center of the simulation voxel.

It is important to note that given the fact that our model assumes only a linear behavior of the vessel (i.e. vessels mimicked only as a resistance component), the expected hemodynamic response function (HRF) characteristics, such as the onset and the undershoot, do not show any modification in the simulated BOLD signal responses. Physiological phenomena such as blood pooling in large venous structures can be modeled by incorporating capacitance elements to the electrical circuit analogy, i.e. an extension of the current model. This addition can then allow variable onsets and post-stimulus undershoot behavior in the simulated BOLD response when caused by delayed blood volume increases. This improvement will help, for instance, to understand the hemodynamic changes across the different cortical depths and the nature of the characteristics of the BOLD HRF.

Further, our approach can be applied to simulate several physiological parameters such as variable blood oxygenation levels, or pathological conditions. Besides, the hemodynamic simulation can be used to investigate the hemodynamic point spread function across the cortical depth and the interpretation of laminar BOLD fMRI (Siero et al, 2011; Fracasso et al, 2018).

### 4.3 On the computation of a dynamic BOLD fMRI signal response

We have simulated the evolution of the BOLD fMRI response using a statistical 3D model of the human cortical vasculature and a hemodynamic response resulting from an arterial dilation as a response to a neuronal drive. This sort of model allows to convert the blood oxygenation changes into spatially dependent magnetic field variations.

Due to the fact that human and mouse cerebral vasculature are similar only to a certain degree, we have found that the frequency components that constitute the evolution of the MR signal for specific oxygenation values are quantitatively different (∼21% difference in frequency components). This demonstrates the impact of the macrovascular architecture between humans and mice on the formation of the BOLD signal.

Moreover, the angular orientation of the vasculature in humans with respect to the main magnetic field strongly modulates the amplitude of the BOLD signal response as demonstrated previously (Fracasso et al, 2018; Viessmann et al, 2019). It is worth to note that pial vessels, according to our simulations, do not directly contribute to the angular dependence seen in previous experimental data (see Figure 7), while intracortical vessels produce the signature cos^2^(φ) modulating behavior in the BOLD signal amplitude. The intracortical cos^2^(φ) modulating behavior might be the result from the elimination of the actual cortical surface in the segmentation process in order to obtain a gray-matter mask and/or from exclusion of pial vessel contributions.

In addition, simulation of water diffusion through these local magnetic field variations provides insights to disentangle quantitative hemodynamic features of the BOLD fMRI signal. The simulated BOLD fMRI signals show qualitative differences in the amplitude for a gradient-echo and spin-echo pulse sequence. This results from the rephasing effect of the RF pulse in the spin-echo, which reduces the effect produced by the macrovascular compartment and highlights the contribution of the microvasculature to the evolution of the BOLD signal (Boxerman et al, 1995).

Due to the linear treatment of the vascular network, i.e. mimicked as a resistance, the onset and post-stimulus undershoot of the BOLD fMRI signal do not present any modification in the simulated acquisition sequence. As mentioned above, physiological phenomena like blood pooling in large venous structures can be modeled by incorporating capacitance elements to the electrical circuit analog. This addition can then demonstrate a different behavior in the simulated BOLD fMRI response when caused by delayed blood volume changes.

The computation of the BOLD fMRI signal assumes only the contribution of the extravascular space, given by the interaction of the diffusing spins and the susceptibility effects induced by the vascular compartments and the differential oxygenation level. The intravascular BOLD fMRI contribution is neglected due to the relatively short T2 relaxation time constant of oxygenated blood at ultra-high magnetic fields (Uludag et al, 2009).

An advantage of our model is the possibility to analyze the laminar BOLD fMRI signal profile dependent on a particular hemodynamic behavior, i.e. laminar flow dynamics and dependencies across the cortical depth. Previous work (e.g. Markuerkiaga et al 2016), has thus far modeled a 2D representation of the vasculature and a static BOLD response across the cortex. The proposed 3D vascular model can improve our understanding of the dynamics in laminar BOLD fMRI signal profiles by enabling simulations of different cortical regions and hemodynamics for different number of arteries and veins (i.e. several input and output sources).

### 4.4 Methodological considerations

The computational pipeline of a representative 3D vascular model, hemodynamics and biophysical effects offers a powerful tool to investigate and disentangle the underlying characteristics of the hemodynamic fingerprint of the BOLD signal. However, our computational pipeline is constrained by assumptions to reduce mathematical complexity and computational load. For instance, the mathematical transcription of the adjacency matrix into an electric circuit is straightforward based on concepts of graph theory. However, the implementation of a non-linear component to simulate a blood reservoir and blood volume changes and the computation hemodynamics become a cumbersome task to resolve.

The simulation pipeline is highly computational time-consuming. For all the described parameters and the cluster server used, significant results are obtained after ∼128 hours of computation. However, the computational time can be reduced with cluster servers that offer better and powerful features or the use of GPU computations.

Moreover, to facilitate investigations on the laminar contributions to the BOLD signal, we also need to consider the difference in cortical thickness between mice and humans. We envision a future improvement of the presented computational model where these limitations will be overcome. For instance, we consider incorporating an active blood flow control on the microvasculature, spatially dependent neuronal activation, and blood volume changes. This improvement will help to understand the hemodynamic changes across cortical depth.

In addition, improvements on the simulation pipeline will include the intravascular contribution that might affect the hemodynamic fingerprint of the BOLD signal evolution. These improvements will inform about the role of the intravascular T2 relaxation time of the blood that depends on magnetic field strength, oxygenation level and applied MR pulse sequence (Grgac et al, 2017; Blockley et al, 2008).

Another important issue to tackle is the validation of our model and results, which of course is an important next step to probe the reliability of our computational model and the accuracy of the simulated BOLD signals as compared to those obtained from the human brain.

## 5. CONCLUSION

We have developed a statistically defined 3D vascular model that resembles the geometrical, topological and rheological characteristics of the human cortical vasculature, and mitigates the current unavailability of a realistic 3D vascular model of the human brain. The proposed computational model permits to simulate in a 3D fashion the hemodynamic changes triggered by a neuronal activation and the local magnetic field disturbances created by the vascular topology and its orientation with respect to the cortical surface and the blood oxygenation changes.

All these interactions provide a comprehensive view on the relation between the vascular topology of the human brain and the associated hemodynamics that result into a *dynamic* BOLD fMRI signal response. This computational pipeline can help the interpretation of the BOLD fMRI signal and can provide a better insight on the quantification of the hemodynamic fingerprint of the signal evolution.

## ACKNOWLEDGEMENTS

This work was supported by the National Institute of Mental Health of the National Institutes of Health under the Award Number R01MH111417

## AUTHOR CONTRIBUTION STATEMENT

MGBY wrote all software, run the simulation pipeline and wrote the original draft of the manuscript. MGBY, JS and NP reviewed and edited the manuscript. All authors discussed the simulation results.

